# A Probabilistic and Anisotropic Failure Metric for Ascending Thoracic Aortic Aneurysm Risk Stratification

**DOI:** 10.1101/2020.09.28.317255

**Authors:** Minliang Liu, Liang Liang, Qing Zou, Yasmeen Ismail, Xiaoying Lou, Glen Iannucci, Edward P. Chen, Bradley G. Leshnower, John A. Elefteriades, Wei Sun

## Abstract

Experimental studies have shown that aortic wall tensile strengths in circumferential and longitudinal directions are different (i.e., anisotropic), and vary significantly among patients with aortic aneurysm. To assess aneurysm rupture and dissection risk, material failure metric of the aortic wall needs to be accurately defined and determined. Previously such risk assessment methods have largely relied on deterministic or isotropic failure metric. In this study, we develop a novel probabilistic and anisotropic failure metric for risk stratification of ascending thoracic aortic aneurysm (ATAA). To this end, uniaxial tensile tests were performed using aortic tissue samples of 84 ATAA patients, from which a joint probability distribution of the anisotropic wall strengths was obtained. Next, the anisotropic failure probability (FP) based on the Tsai−Hill (TH) failure criterion was derived. The novel FP metric, which incorporates uncertainty in the anisotropic failure properties, can be evaluated after the aortic wall stresses are computed from patient-specific biomechanical analysis. For method validation, “ground-truth” risks of additional 41 ATAA patients were numerically-reconstructed using corresponding CT images and tissue testing data. Performance of different risk stratification methods (e.g., with and without patient-specific hyperelastic properties) was compared using p-value and receiver operating characteristic (ROC) curve. The results show that: (1) the probabilistic FP metric outperforms the deterministic TH metric; and (2) patient-specific hyperelastic properties can help to improve the performance of probabilistic FP metric in ATAA risk stratification.

## 1. Introduction

Thoracic aortic aneurysm is a lethal disease, which may lead to aortic rupture or dissection: the five-year survival in patients left untreated is 54% [1]. Currently, the clinical surgery criterion is primarily based on the aortic size and classifies an ascending thoracic aortic aneurysm (ATAA) as high risk if the (maximum) diameter is larger than 5.5cm [2, 3], which may not accurately reflect a patient’s risk [4, 5]: some aneurysms at smaller diameters (e.g., < 4cm) can and do rupture [4]. As ATAA rupture and dissection are essentially mechanical events, patient-specific biomechanical assessment, such as structural finite element analysis (FEA), can provide a more accurate assessment of ATAA rupture/dissection risk [6, 7]. Using 3D patient-specific aorta geometry and physiological blood pressure, aortic wall stresses can be computed using FEA. To assess patient-specific ATAA rupture/dissection risk, computed stress distributions on the aortic wall at physiological, hypertensive, or elevated blood pressures are compared with the tensile strengths of the aortic wall, by which a scalar-valued failure metric can be obtained. Therefore, an accurate failure metric plays a critical role in biomechanical ATAA risk assessment [8].

Isotropic material failure metrics, such as the von Mises stress equivalent stress [9-11], have been widely adopted in biomechanical risk assessment of aortic aneurysms. Another popular failure metric is the rupture potential index (RPI) [12], which is obtained by dividing an isotropic wall stress (e.g., maximum principal stress [13]) by wall strength. These isotropic criteria may not be appropriate for aortic tissues [14], because the aortic wall strengths in the circumferential and axial directions are significantly different, which has been revealed by many experimental studies [15-20]. To incorporate direction-dependent failure properties, anisotropic failure metric needs to be used. The Tsai−Hill (TH) criterion [21] is a well-known anisotropic failure model that was originally developed for engineered fiber-reinforced composites. Recently, Korenczuket al. [14] applied the TH failure metric to characterize the failure of porcine abdominal aortas, and the authors found that the anisotropic TH metric performed much better than the isotropic von Mises stress metric [14].

To perform patient-specific biomechanical assessment, parameters of a failure metric (e.g., wall strength in the RPI) need to be quantified. Some studies suggested to use deterministic approaches [13, 22-24]. For instance, Geest et al. [25] proposed a linear regression model to estimate wall strength of abdominal aortic aneurysm (AAA) from patient parameters (age, gender, maximum dimeter, family history and smoking status) and local parameters (local intraluminal thrombus (ILT) thickness and local diameter). However, large variability in aortic wall strength has been revealed by many experimental works [15, 17-19], which indicate that the predictive capability of the linear regression model is limited [25]. Indeed, patient-specific failure properties (i.e., aortic wall strengths) can only be accurately determined using invasive and destructive tests, and these tests clearly cannot be performed for patients whose ATAAs are still intact. Recently, a probabilistic rupture risk index (PRRI) [26] was proposed for biomechanical risk assessment of AAA. The PRRI incorporates a probability distribution of the wall strength, and it offers a physical meaning: PRRI represents the probability of failure. The PRRI outperformed the deterministic RPI in a retrospective study [7] of asymptomatic AAA. However, the PRRI was developed based on the isotropic maximum principal stress, ignoring the fact that the wall strengths are direction dependent.

In this work, we developed a novel probabilistic and anisotropic failure metric for ATAA risk stratification. Uniaxial tensile tests were performed using aortic tissue samples from 84 ATAA patients, from which a two-dimensional (2D) probability distribution of the anisotropic wall strengths was obtained. Next, the anisotropic failure probability (FP) based on the TH failure theory was derived from the probability distribution of wall strength. After the aortic wall stresses are computed from patient-specific biomechanical analyses, the novel FP metric can be used for risk classification. For validation, the “ground-truth” risks of additional 41 ATAA patients were reconstructed from FE simulations using the CT images and tissue testing data (planar biaxial for hyperelastic properties and uniaxial for failure properties) of the 41 patients. Using the “ground-truth” data, different risk stratification methods and failure metrics were compared.

## 2 Methods

### 2.1 Aortic tissue samples and CT image data

In this study, surgically resected human aortic tissues of 98 ascending thoracic aortic aneurysm (ATAA) patients (72 males, 26 females, age: 62.57 ± 12.42 years) were obtained from the Emory Saint Joseph’s Hospital with IRB approval. The tissue samples underwent uniaxial tests (Section 2.2) to determine the failure properties, and biaxial tests (Section 2.4) to determine hyperelastic properties. Among the 98 patients, pre-operative 3D CT images of 14 patients were obtained to reconstruct their ATAA geometries. In addition, we obtained surgically resected aortic tissue samples and corresponding CT images of 27 ATAA patients in a previous study [27] from Yale-New Haven Hospital. Uniaxial and biaxial testing were performed for the 27 patients.

The cohort of 125 (98+27) ATAA patients was divided into two groups: (1) 84 patients without CT images; and (2) 41 patients with CT images (including 27 patients from our previous study [27]). Group 1 was used for method development (Section 2.3); and Group 2 was used to compute the “ground-truth” risks for method validation (Section 2.5). Hence, there is no overlapping between the two groups. Patient characteristics in Group 2 are reported in Table 1.

**Table 1.** Patient characteristics in Group 2 used for validation. The reconstructed failure pressure is obtained in Section 2.5.

| Patient # | Age | Gender | Diameter (mm) | Systolic pressure ($P_{sys}$, mmHg) | Reconstructed failure pressure ($P_f$, mmHg) |
| --- | --- | --- | --- | --- | --- |
| 1 | 76 | M | 46 | 130 | 442 |
| 2 | 52 | M | 47 | 101 | 426 |
| 3 | 69 | F | 47 | 150 | 461 |
| 4 | 66 | M | 46 | 131 | 252 |
| 5 | 73 | M | 40 | 147 | 658 |
| 6 | 73 | M | 47 | 178 | 222 |
| 7 | 69 | F | 50 | 138 | 397 |
| 8 | 59 | M | 51 | 128 | 239 |
| 9 | 60 | M | 48 | 128 | 453 |
| 10 | 59 | M | 47 | 119 | 151 |
| 11 | 45 | M | 54 | 144 | 336 |
| 12 | 48 | M | 52 | 119 | 191 |
| 13 | 48 | M | 51 | 105 | 403 |
| 14 | 59 | M | 53 | 131 | 414 |
| 15 | 64 | F | 49 | 104 | 280 |
| 16 | 62 | F | 50 | 75 | 200 |
| 17 | 66 | M | 55 | 134 | 272 |
| 18 | 52 | M | 48 | 133 | 201 |
| 19 | 56 | M | 50 | 128 | 208 |
| 20 | 47 | M | 53 | 97 | 202 |
| 21 | 56 | M | 46 | 139 | 191 |
| 22 | 57 | M | 52 | 136 | 428 |
| 23 | 71 | M | 48 | 105 | 242 |
| 24 | 33 | M | 41 | 136 | 145 |
| 25 | 44 | M | 36 | 155 | 408 |
| 26 | 56 | F | 58 | 117 | 262 |
| 27 | 68 | M | 52 | 123 | 295 |
| 28 | 65 | M | 50 | 138 | 377 |
| 29 | 46 | F | 57 | 135 | 190 |
| 30 | 55 | M | 52 | 129 | 191 |
| 31 | 57 | M | 63 | 132 | 153 |
| 32 | 58 | M | 43 | 137 | 267 |
| 33 | 58 | M | 48 | 142 | 316 |
| 34 | 68 | M | 47 | 139 | 214 |
| 35 | 70 | M | 66 | 124 | 141 |
| 36 | 65 | M | 51 | 139 | 320 |
| 37 | 44 | M | 56 | 153 | 302 |
| 38 | 77 | F | 51 | 130 | 195 |
| 39 | 56 | M | 49 | 124 | 379 |
| 40 | 31 | F | 57 | 93 | 274 |
| 41 | 68 | M | 46 | 146 | 286 |

### 2.2 Uniaxial Tests and Failure Criterion

The human ATAA samples were cryopreserved in 90%/10% RPMI-1640/DMSO solution and stored at -80□°C until testing. Frozen ATAA samples were slowly defrosted, followed by the multi-stage slow thawing process to remove the cryopreservation agent [15, 28]. Uniaxial tensile tests were then performed using the 98 ATAA tissue samples from Emory (similar uniaxial tests were performed for the 27 patients in our previous study [27]). The ATAA samples were trimmed into 25×13 mm dog bone-shaped specimens using a 3D printed template. For each ATAA patient, circumferential and axial specimens were obtained and subjected to uniaxial tensile tests until failure, from which circumferential and axial strengths were extracted. Thickness was measured at three locations in the narrow portion for each specimen with a Mitutoyo 7301 rotating thickness gage (Aurora, IL) that has an accuracy of ±0.01 mm. An average thickness was determined from the three measurements and used in the calculation of the undeformed cross-sectional area. The uniaxial tests were conducted at room temperature using a Test Resources 100Q Universal Testing Machine (Shakopee, MN). Graphite markers were placed on the narrow portion of the specimens for optical strain measurements. The axial force was measured using a 4.4N load cell (Test Resources SM-500-294) [29]. Fine grit sandpaper was placed between the tissue and the clamps to avoid slippage during the test. The specimens were continuously hydrated with 0.9% saline solution during testing [30]. The specimens were quasi-statically stretched to failure at a constant displacement rate of 5mm/min.

The testing data were excluded from subsequent analyses if rupture did not occur in the middle region of the test specimen [20, 31]. We observed that the aortic wall layers, i.e., media and adventitia layers, may rupture at different times during testing. Therefore, failure was defined by the onset of yielding, i.e., a deviation from the exponential strain-stress curve, which corresponds to a rupture of any layer. Results of the uniaxial testing are summarized in Figure 1(a) for the two groups described in Section 2.1.

**Figure 1.**
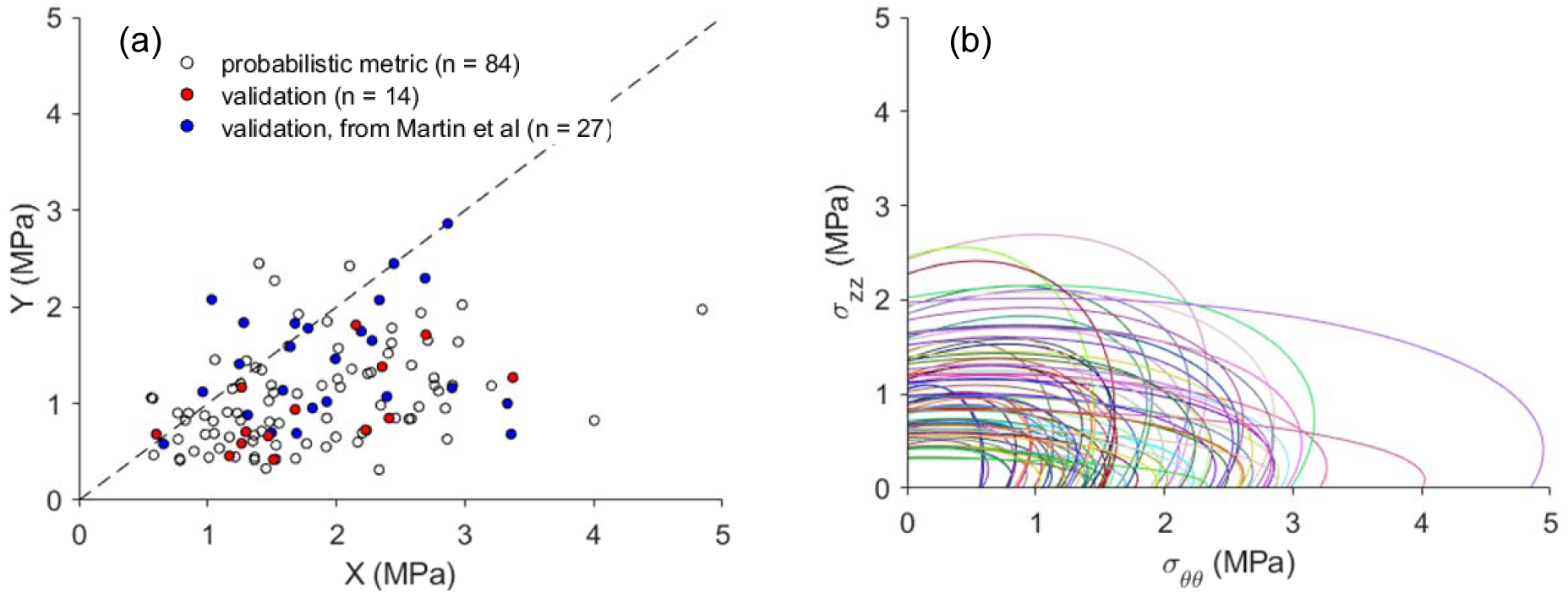
Uniaxial strengths and failure envelopes. (a) Circumferential (*X*) and axial (*Y*) strengths of the 125 ATAA patients. Among them, 84 patients were used for developing the probabilistic metric (Group 1), 41 patients were used for validation (Group 2). (b) Tsai−Hill (TH) failure envelopes of the 84 patients in Group 1. *τ*_*θz*_ = 0 when generating the failure envelopes.

In this study, the Tsai−Hill (TH) criterion [21] was used for modeling failure properties of the aortic tissues. The TH model has demonstrated a good fitting capability with off-axis tension test data of aortic tissues [14, 20]. Applying the TH criterion to the aortic tissue, the failure metric takes the following form:

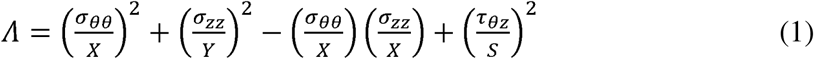

where *σ*_*θθ*_, *σ*_*ZZ*_ and *τ*_*θZ*_ stand for circumferential stress, axial stress and in-plane shear stress, respectively. Cauchy stress is used when the TH criterion is applied for finite deformation [20]. *X,Y* and *S* are circumferential, axial and in-plane shear strengths, respectively, which are the model parameters to be determined. Failure happens when *Λ* reaches 1.

Due to the limited size of tissue samples, we were unable to perform off-axis uniaxial tests to obtain. An average value (*S* = 0.635*MPa*) from our previous study [20] is used for all patients. The original TH model [21] assumes *X >Y*, which is usually the case for unidirectional fiber-reinforced composite. For the tested aortic tissue samples, *X < Y* can be observed for a few patents (see Figure 1(a)). Therefore, in this study, the TH failure metric is modified as

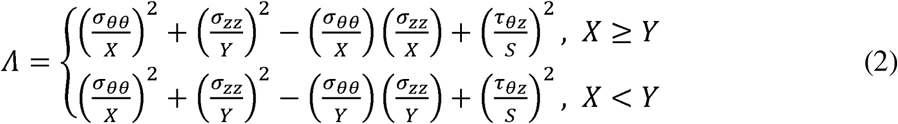

Hence, using the circumferential (*X*) and axial (*Y*) strengths determined from uniaxial tests, the TH failure envelopes (contour line defined by *Λ* = 1) of the 84 patients in Group 1 can be visualized in Figure 1(b) using *τ*_*θz*_ = 0.

### 2.3 Anisotropic Probabilistic Failure Metric

A anisotropic probabilistic failure metric is developed in following the steps: (1) estimating the probability density function (PDF) of failure parameters *X* and *Y* (circumferential and axial strengths), *f*_*XY*_, from uniaxial testing data of Group 1; (2) deriving PDF of the failure metric *Λ, f*_*Λ*_, by using the method of random variable transformation (a.k.a. change of variables technique) [32]; and (3) calculating the failure probability (FP), *P(*Λ* >* 1).

In this study, the joint PDF *f*_*XY*_ of the TH model parameters *X* and *Y* (circumferential and axial strengths) is obtained using kernel density estimation (KDE) [33, 34], which is a non-parametric model for PDF estimation. The bivariate KDE [35] with diagonal bandwidth matrix takes the following form

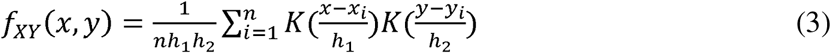

where *x*_*i*_ and *y*_*i*_ are samples of the TH model parameters *X* and *Y*.*n* is the number of samples. *h*_1_ and *h*_2_ represent the bandwidths. *K*(*▪*) is the one-dimensional kernel function. In this study, the normal kernel was selected:

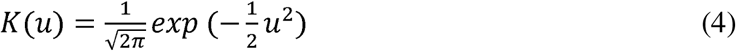

To ensure that the support for the PDF is positive, i.e.,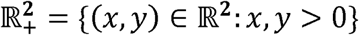, log transformation was used [36]. *h*_1_ and *h*_2_ were chosen using ten-fold cross validation with likelihood function of the 84 patients data (Group 1) in following the steps: (1) *h*_1_ and *h*_2_ are evenly sampled on a 2D grid from 0.1 to 0.35 with a total of 10,000 points; (2) using a set of *h*_1_ and *h*_2_ values, ten-fold cross validation is performed to evaluate the log-likelihood function; (3) the *h*_1_ and *h*_2_ that lead to the maximum log-likelihood in the ten-fold cross validation were chosen. As a result, *h*_1_ *=* 0.2439*MPa* and *h*_2_ *=* 0.2439*MPa*. In Figure 2, the joint PDF *f*_*XY*_ and marginal PDFs *f*_*x*_ and *f*_*Y*_ are plotted.

**Figure 2.**
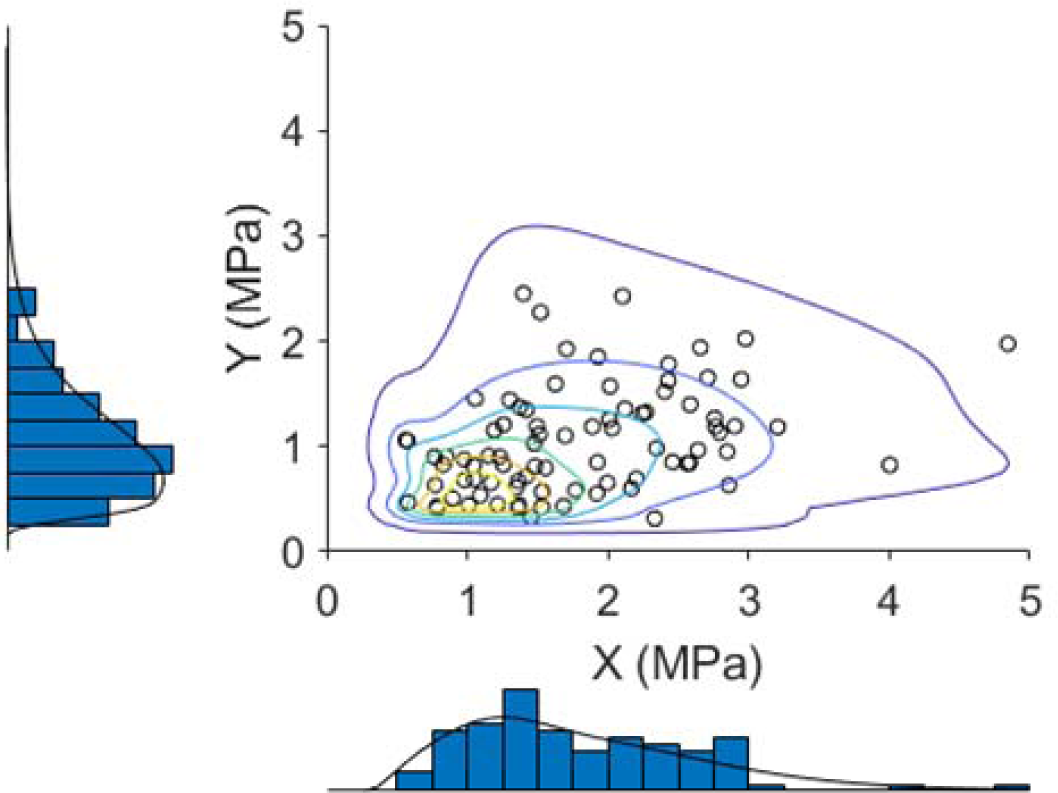
The KDE-estimated joint PDF *f*_*XY*_ and marginal PDFs *f*_*x*_ and *f*_*Y*_. Histograms for marginal distributions are also visualized.

To validate the KDE-estimated distribution, goodness-of-fit tests were performed using the estimated joint PDF *f*_*XY*_ and marginal PDFs. The results are summarized in Table 2, *p* =0.9715 was obtained using Chi-square goodness-of-fit test for the joint distribution. The null hypothesis is that the data comes from the KDE-estimated distribution. For all tests, the null hypothesis cannot be rejected, which indicates the PDFs can well describe the data distribution.

**Table 2.** Goodness-of-fit tests for the joint distribution and marginal distributions. Null hypothesis: the data comes from the KDE-estimated distribution; alternative hypothesis: the data does not come from such distribution.

| type of test | joint PDF $f_{XY}$ | type of test | marginal PDF $f_X$ | marginal PDF $f_Y$ |
| --- | --- | --- | --- | --- |
| Chi-square | 0.9715 | Chi-square | 0.6481 | 0.9449 |
|  |  | Kolmogorov-Smirnov | 0.5857 | 0.9346 |
|  |  | Anderson-Darling | 0.6432 | 0.9030 |

To derive the PDF *f*_*Λ*_ of the TH failure metric *Λ*, the method of direct transformation [32] is employed. Given the stress states (*σ*_*θθ*_, *σ*_*ZZ*_ and *τ*_*θz*_), is a function of the random variables, i.e., *Λ* = *Λ* (*X, Y*). Since *S* is a fixed constant in this study, we define a random variable *W*,

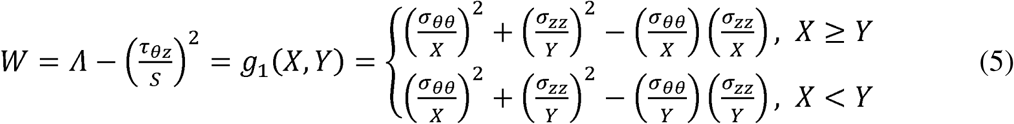

Therefore, the probability of failure *P*(*Λ >* 1) is equivalent to 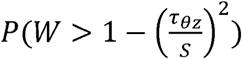. We also define a dummy random variable *Z*

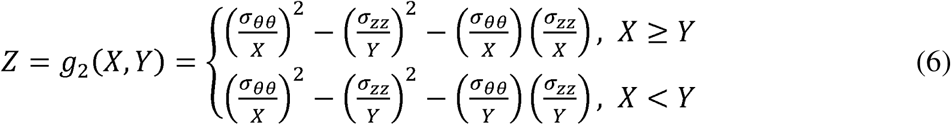

The two functions *g*_1_(*X,Y*) and *g*_2_(*X,Y*) transform *X* and *Y* to the space of *W* and *Z*. Let 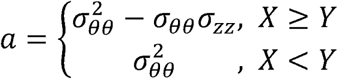, and 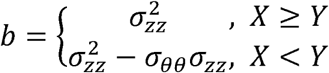, then we can rewrite the transformation as 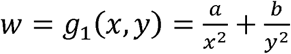, and 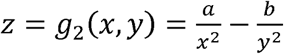, with support 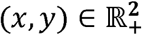 Hence, the inverse transformation and its Jacobian can be obtained, i.e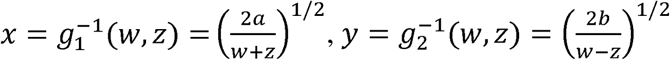 ^and^ 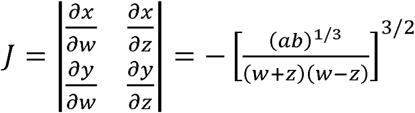.

When *a* ≠ 0 and *b* ≠0, the transformation from (*X, Y*) to (*W, Z*) is one-to-one because j ≠ 0. Using the method of direct transformation [32], the joint PDF of *W* and *Z, f*_*WZ*_, can be derived using

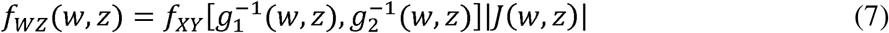

The marginal PDF *f*_*W*_ can be obtained by

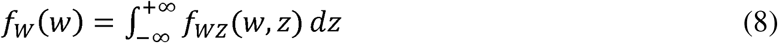

When *a* = 0 or *b* = 0, *f*_*w*_ can be directly obtained using the method of direct transformation for one random variable [32]. If *a* = 0, i.e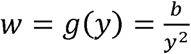, PDF *f*_*w*_ can be derived as

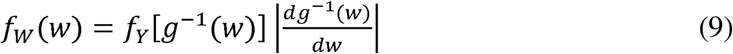

Similarly, if *b* = 0, i.e., 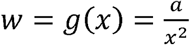, *f*_*w*_ can be obtained by

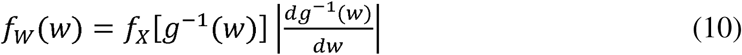

For illustrative purposes, *f*_*w*_ is plotted in Figure 3 by using different values of *σ*_*θθ*_ and *σ*_*zz*_. Once the PDF *f*_*w*_ is obtained, it is straightforward to compute the failure probability (FP) by integration:

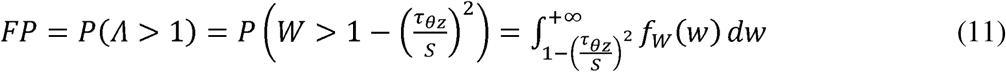

**Figure 3.**
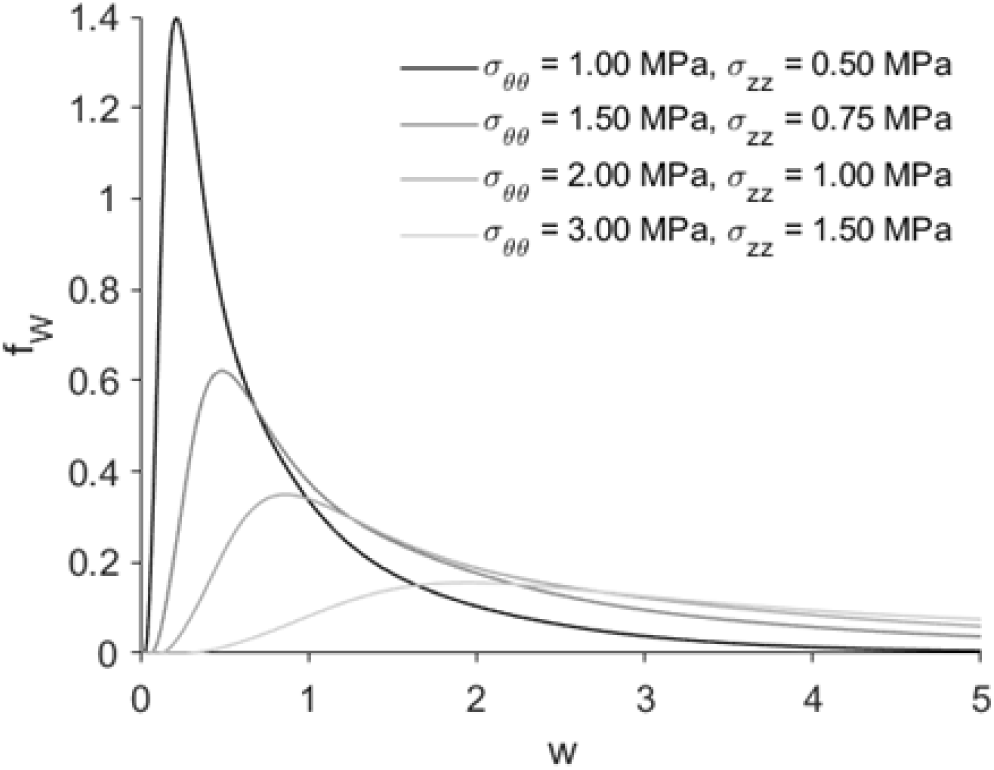
The estimated pdf *f*_*w*_ with given *σ*_*θθ*_ and *σ*_*zz*_ values.

Given *τ*_*θz*_, FP can be visualized on a 2D plot of *σ*_*θθ*_ and *σ*_*zz*_. In Figure 4, 2D contour plots of FP are generated using representative *τ*_*θz*_ values.

**Figure 4.**
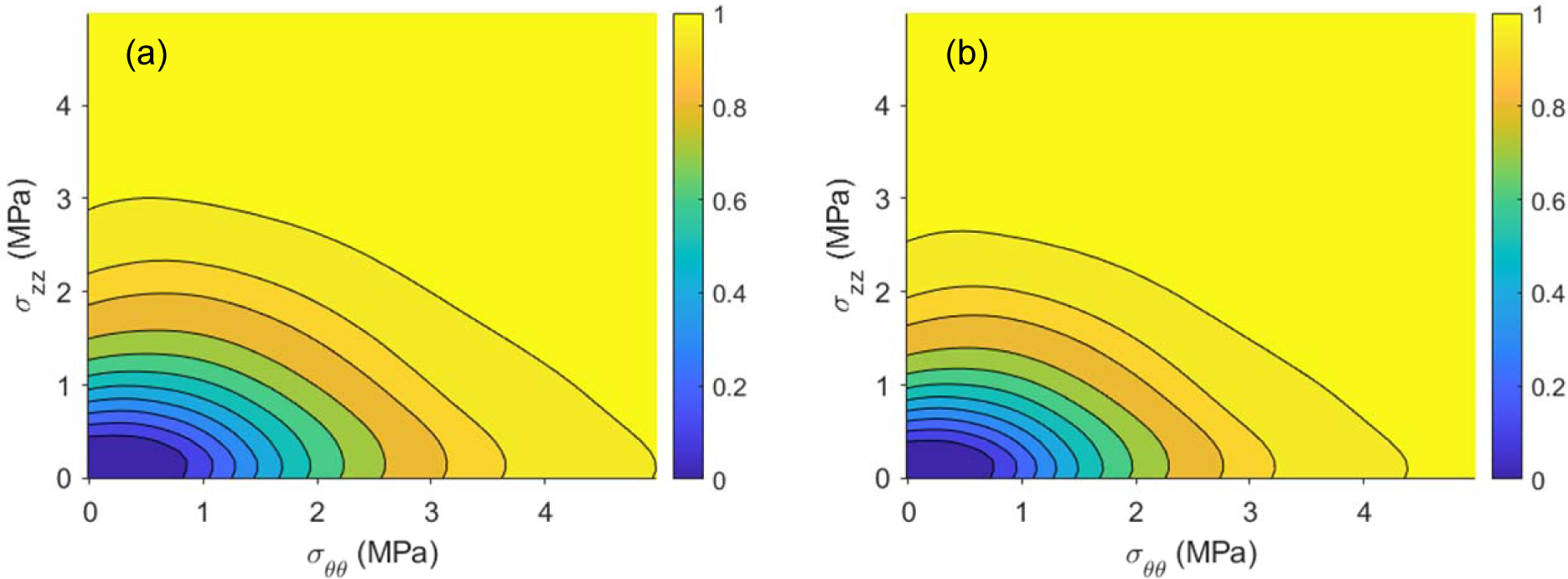
2D contour of FP in *σ*_*θθ*_ *— σ*_*zz*_ plane using (a) *τ*_*θz*_ *=* 0 and (b) *τ*_*θz*_ = *300kPa*.

Given stress states, the value of FP can be obtained via numerical integration. In this study, the FP values were pre-computed in a 3D mesh grid of *σ*_*θθ*_, *σ*_*zz*_ and *τ*_*θz*_ prior to FE simulations. FP under specified stress states can be obtained using 3D interpolations from the pre-computed FP values in the grid.

### 2.4 Biaxial Tests and Hyperelsatic Model

Seven-protocol biaxial tensile tests were performed on the patient tissues in Group 2. Since the test results for the 27 Yale patients were already available from our previous study [27], in this study, we only performed the tests on the tissues of the 14 patients from Emory. Briefly, the samples were carefully trimmed into square-shaped specimens with a side length of 20∼25 mm. Thickness was measured at three locations on the diagonal line of the biaxial specimen. An averaged undeformed thickness was recorded. Each specimen was subjected to biaxial tensile loadings which aligned with the circumferential and axial directions. A stress-controlled biaxial testing protocol was utilized [15]. *N* denotes the nominal stress, and the ratio ***N***_*θθ*_:*N*_*zz*_ was kept constant. Each tissue specimen was preconditioned for at least 40 continuous cycles with *N*_*θθ*_:*N*_*zz*_ = 1:1 to minimize hysteresis. Seven successive protocols were performed using ratios *N*_*θθ*_:*N*_*zz*_ = 0.3:1, 0.5:1, 0.75:1, 1:1, 1: 0.75, 1: 0.5, 1: 0.3.

To describe the hyperelastic properties of the aortic wall, the Gasser-Ogden-Holzapfel (GOH) model [37] was used. In the GOH model, tissues are assumed to be composed of an isotropic matrix with two families of fibers, each of which has a preferred direction. The fiber dispersion is modeled using a generalized structural tensor (GST) approach. The strain energy density function can be expressed by

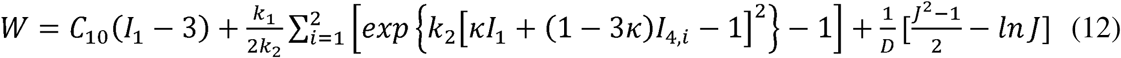

where *C*_10_ is a material parameter to describe the isotropic matrix. *k*_1_ is a positive material parameter that has the same unit of stress, while *k*_2_ is a dimensionless parameter. *I*_1_ = *tr(**C**)* is the first strain invariant, where ***C* = *F***^***T***^***F. I***_***4i***_ **= *a***_***0i***_ •(***Ca***_***0i***_***), i* =** 1,2 are two additional pseudoinvariants that describe deformations in the preferred fiber directions, in which unit vectors ***a***_01_ and ***a***_02_ characterize the two fiber directions in the reference configuration. *θ* is the angle between a fiber direction and the axis of symmetry (circumferential direction of the aortic wall). For instance, ***a***_01_ = (*cos θ, sin θ*,0) and ***a***_02_ = (*cos θ, – sin* θ,0). *k* is used as a dispersion parameter describing the distribution of fiber orientations. When *k* = 0, the fibers are perfectly aligned. When *k* = 0.33, the fibers are uniformly distributed, and the material becomes isotropic. *D* is a fixed constant enforcing the material incompressibility (*D* = 1 × 10^−5^). The five material parameters (*C*_10_, *k*_1_ *k*_2_, *k, θ*) for an ATAA patient were determined by fitting the GOH model to the biaxial data.

### 2.5 Reconstructing ATAA Risk using Patient-Specific CT Images and Tissue Testing Data

To reconstruct ATAA risk of adverse events (dissection or rupture) for the 41 patients in Group 2, FE simulations were performed using patient-specific information (geometry, blood pressure, experimentally-measured wall thickness, hyperelastic parameters and failure parameters) in Abaqus (Figure 5). The patient-specific ATAA geometries were segmented from the patients’ pre-operative CT scans using 3D Slicer [38], we assume these geometries are in equilibrium under the systolic blood pressure. Then, the ATAA surfaces were meshed into quadrilateral elements using our remeshing program [39]. Mesh sensitivity analysis was performed in our previous study [27]. To estimate the patient-specific wall thickness in the systolic phase, an analytical procedure [40] was performed using the experimentally-measured undeformed wall thickness assuming *σ*_*θθ*_: *σ*_*zz*_ = 1:2 [40]. Solid meshes of the ATAA systolic geometries with C3D8H elements in Abaqus were then obtained by extruding the surface mesh using corresponding systolic wall thickness values.

**Figure 5.**
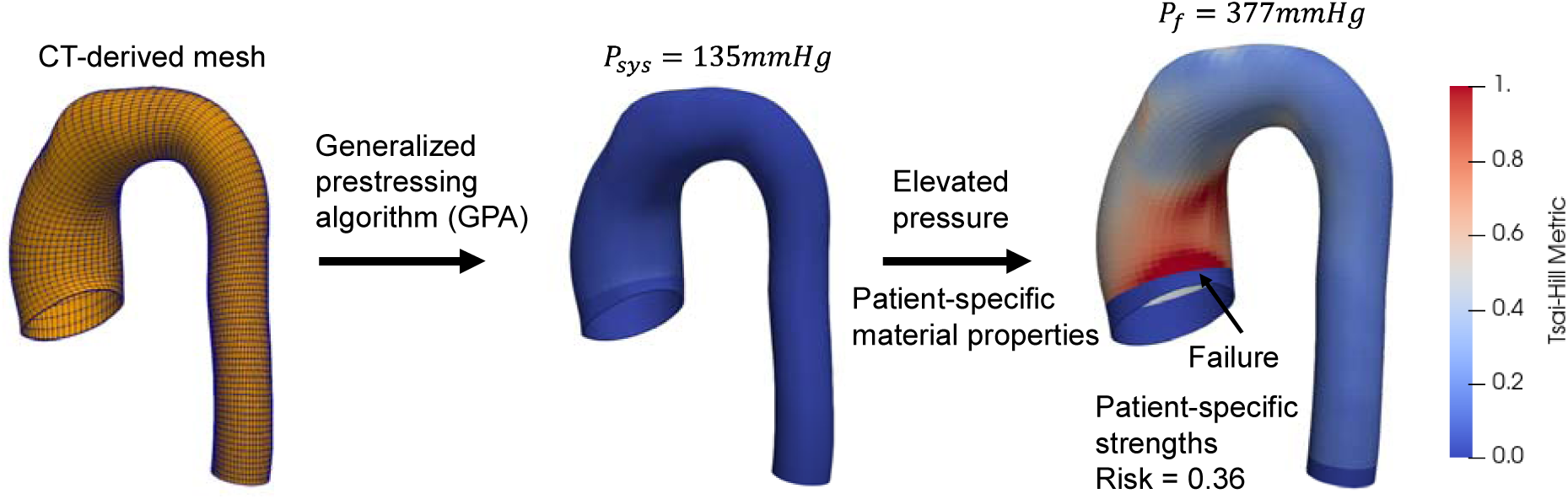
FE simulations to reconstruct ATAA risk.

**Figure 6.**
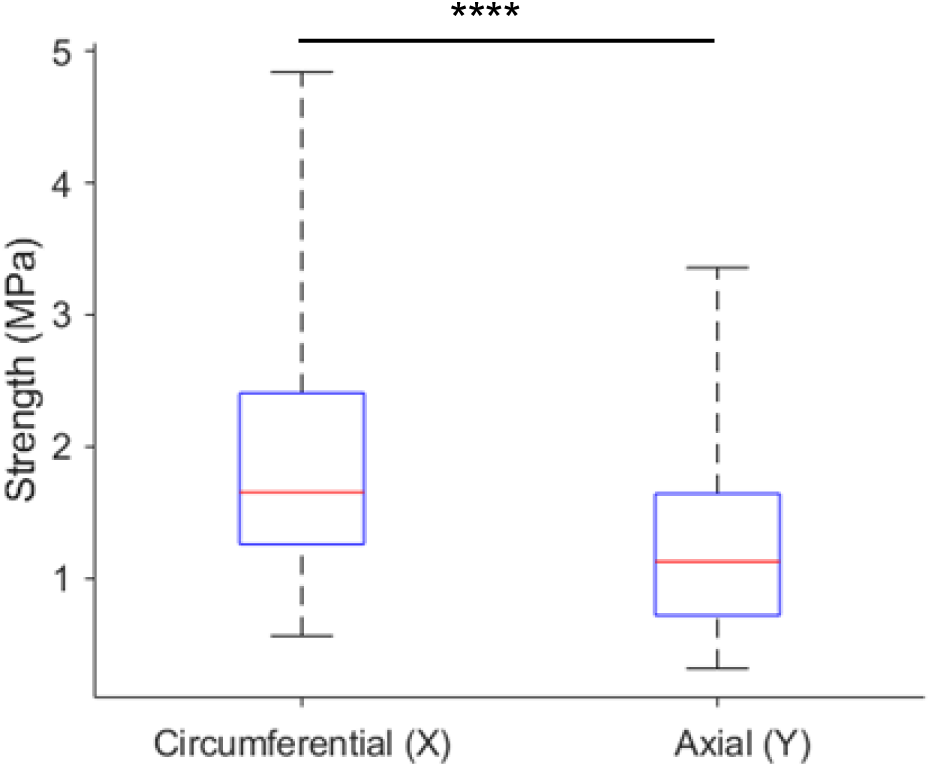
Boxplot of uniaxial strengths of human ATAA tissues. The red mark indicates the median, and the bottom and top edges of the box indicate the 25th and 75th percentiles, respectively. The whiskers extend to the maximum and minimum. **** indicates statistical significance level *p* ≤ 0.0001.

The generalized pre-stressing algorithm (GPA) [41] was utilized to incorporate the prestress induced by the systolic pressure into the patient-specific ATAA systolic geometries. In GPA, the total deformation gradient ***F***_*t*_ is stored as a history variable for each integration point. The ***F***_*t*_ is updated based on the incremental deformation gradient Δ***F*** resulting from the prescribed loading and boundary conditions.

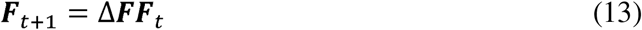

The incremental deformation gradient Δ***F*** resulting from the systolic pressure is iteratively applied to the systolic geometries and stored in ***F***_*t*_.

It is observed that ATAA dissection and rupture usually occurs under elevated blood pressures brought on by extreme emotional or physical stress [42, 43]. To this end, using the patient-specific GOH material parameters, the aorta geometries are pressurized under elevated pressure levels until failure is predicted by the patient-specific TH parameters measured from uniaxial tests (see Figure 5). In the FE simulations, the proximal and distal boundaries of the ATAA geometries were only allowed to move in the radial direction. To remove boundary effect, three layers of elements adjacent to the boundaries were excluded from the failure evaluation. For a particular patient, the pressure rupture risk (PRR) [40] is used to quantify the risk revealed by the FE simulation, which is defined as

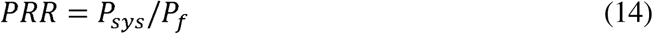

where *P*_*sys*_ stands for the patient’s systolic blood pressure and *P*_*f*_ represents the patient’s failure pressure. We define high risk as for patient who has a *PRR* ≥ 0.6 and low risk as *PRR <* 0.6. According to American Heart Association (AHA)’s blood pressure category [44], if *P*_*sys*_ is normal, *Pf* = *P*_*sys*_*/0*.*6* corresponds to the hypertensive crisis (blood pressure higher than 180mmHg). Therefore, using *PRR* = 0.6 as threshold, high risk patients are prone to adverse events under the hypertensive crisis. The numerically-reconstructed risks are considered as “ground-truth” for evaluation of the risk stratification methods in Section 2.6.

### 2.6 ATAA Risk Stratification Methods

To estimate a patient’s risk, we can only use information (from medical report data or clinical images) before any adverse event, which may include geometries from images and blood pressure. We note that patient-specific failure information (wall strengths and failure pressure) cannot be used in the risk stratification, as those can only be obtained using invasive and destructive testing. Using stresses computed from FE simulation, predefined failure metric can be evaluated on the ATAA wall under a specified pressure for an individual patient. According to several studies, wall thickness may be measured from MRI [45-48] or high resolution CT scans [49]. Hence, in this study, the same 3D aortic geometries from Section 2.5 were used for risk stratification, representing an ideal scenario in which wall thickness can be measured from images.

We apply four different methods to classify high (*PRR* > 0.6) and low risk (*PRR* < 0.6) patients, and method performance is evaluated using p-value and receiver operating characteristic (ROC) curve. The following risk stratification methods are investigated:

1. Maximum diameter criterion/metric.
2. Maximum failure metric at *P*_*sys*_ (hyperelastic properties are not needed).
3. Maximum failure metric at 1.5*P*_*sys*_ using one set of hyperelastic properties representing the population-mean response.
4. Maximum failure metric at 1.5*P*_*sys*_ using patient-specific hyperelastic properties.

The metric in Method 2 to Method 4 is evaluated at a spatial point on the aortic wall. The maximum failure metric is the maximum among the metric values from all spatial points on the aortic wall. By using static determinacy, the transmurally-mean wall stress can be readily obtained from clinical images without knowing the patient-specific hyperelastic properties [50-52]. Here, a set of material parameters in the GOH model defines the hyperelastic properties of a patient’s aortic wall. Therefore, in Method 2, patient-specific hyperelastic properties are not needed to compute maximum failure metric at *P*_*sys*_. Failure metric under elevated blood pressure (e.g., 1.5*P*_*sys*_) may provide more valuable insight on the ATAA risk. However, under supra-physiological pressure loads, failure metric depends on the patient-specific hyperelastic properties. A study [26] suggests to use a population-mean response as a surrogate. Hence, to evaluate this strategy, one representative set of hyperelastic properties is used for all patients in Method 3. Patient-specific hyperelastic properties can be identified from multi-phase clinical images using inverse approaches [53-58]. Therefore, in Method 4, patient-specific hyperelastic properties are used to evaluate the maximum failure metric at 1.5*P*_*sys*_.

For comparisons, the probabilistic metric FP (Eqn. (11)) and a deterministic Tsai−Hill (TH) metric (Λ in Eqn. (2) with typical parameters *X* = 2.5*MPa* and *Y* = 1.2*MPa*) are used as the failure metric in Method 2 and Method 4.

## 3. Results

### 3.1. Circumferential and Axial Strengths

To demonstrate that the wall strength of the human ATAA tissues is anisotropic, circumferential and axial strengths of the 125 ATAA patients are depicted in Figure 5 using boxplot. By using two-sample t-test, it is shown that the circumferential strength is significantly higher than the axial strength with *p <* 0.0001.

### 3.2. Reconstructed ATAA Risk

Using the patient-specific imaging and tissue testing data, failure pressure (*P*_*f*_) and risk (PRR) were numerically-reconstructed for the 41 ATAA patients in Group 2 and considered as “ground-truth” for risk stratification. The reconstructed (*P*_*f*_) is shown in Table 1 and the PRR is listed in Table 3.

**Table 3.** Reconstructed risk (PRR) and failure metrics evaluated by different methods (Section 2.6).

| Patient # | Method 2 (FP) | Method 3 (FP) | Method 4 (FP) | Method 2 (TH) | Method 4 (TH) | Risk (PRR) |
| --- | --- | --- | --- | --- | --- | --- |
| 1 | 0.0054 | 0.0960 | 0.0922 | 0.0519 | 0.1571 | 0.2944 |
| 2 | 0.0436 | 0.1715 | 0.1412 | 0.1048 | 0.2127 | 0.2373 |
| 3 | 0.0311 | 0.1310 | 0.1596 | 0.0888 | 0.2139 | 0.3251 |
| 4 | 0.0385 | 0.2749 | 0.3350 | 0.1012 | 0.3271 | 0.5200 |
| 5 | 0.0010 | 0.0339 | 0.0224 | 0.0329 | 0.0848 | 0.2236 |
| 6 | 0.1708 | 0.4672 | 0.4416 | 0.2030 | 0.4658 | 0.8003 |
| 7 | 0.0022 | 0.0601 | 0.0392 | 0.0434 | 0.0960 | 0.3477 |
| 8 | 0.0112 | 0.1813 | 0.1750 | 0.1054 | 0.2972 | 0.5366 |
| 9 | 0.0159 | 0.1325 | 0.1279 | 0.0625 | 0.1595 | 0.2829 |
| 10 | 0.4105 | 0.2408 | 0.6849 | 0.5012 | 1.0244 | 0.7859 |
| 11 | 0.0956 | 0.3372 | 0.4444 | 0.1371 | 0.4891 | 0.4290 |
| 12 | 0.0371 | 0.0329 | 0.3790 | 0.1317 | 0.4837 | 0.6245 |
| 13 | 0.0104 | 0.0991 | 0.1315 | 0.0671 | 0.1675 | 0.2606 |
| 14 | 0.0047 | 0.1136 | 0.1449 | 0.0510 | 0.1580 | 0.3167 |
| 15 | 0.0001 | 0.0093 | 0.0148 | 0.0213 | 0.0637 | 0.3714 |
| 16 | 0.0038 | 0.0747 | 0.1077 | 0.0500 | 0.1614 | 0.3759 |
| 17 | 0.0208 | 0.2579 | 0.3149 | 0.1001 | 0.3482 | 0.4926 |
| 18 | 0.0251 | 0.2182 | 0.2834 | 0.0729 | 0.2723 | 0.6627 |
| 19 | 0.1310 | 0.4879 | 0.5801 | 0.1748 | 0.6561 | 0.6152 |
| 20 | 0.0058 | 0.1791 | 0.2325 | 0.1183 | 0.3835 | 0.4796 |
| 21 | 0.0470 | 0.2843 | 0.3010 | 0.1044 | 0.2890 | 0.7287 |
| 22 | 0.0100 | 0.1273 | 0.1076 | 0.0657 | 0.1538 | 0.3176 |
| 23 | 0.0910 | 0.2442 | 0.2884 | 0.1557 | 0.3810 | 0.4330 |
| 24 | 0.0745 | 0.5103 | 0.6349 | 0.2220 | 0.8389 | 0.9393 |
| 25 | 0.0116 | 0.0414 | 0.0389 | 0.0626 | 0.0897 | 0.3800 |
| 26 | 0.0115 | 0.1103 | 0.0969 | 0.1065 | 0.2426 | 0.4473 |
| 27 | 0.0044 | 0.0914 | 0.0945 | 0.0510 | 0.1341 | 0.4163 |
| 28 | 0.0254 | 0.2496 | 0.2160 | 0.0828 | 0.2267 | 0.3583 |
| 29 | 0.1929 | 0.0624 | 0.3067 | 0.2352 | 0.3430 | 0.6792 |
| 30 | 0.0320 | 0.2833 | 0.3067 | 0.0876 | 0.3073 | 0.6895 |
| 31 | 0.2592 | 0.6240 | 0.8115 | 0.2884 | 0.8678 | 0.8948 |
| 32 | 0.1280 | 0.4604 | 0.4085 | 0.1487 | 0.3659 | 0.5327 |
| 33 | 0.1949 | 0.4687 | 0.4188 | 0.2125 | 0.4203 | 0.4399 |
| 34 | 0.2138 | 0.0889 | 0.4364 | 0.2870 | 0.5355 | 0.5800 |
| 35 | 0.1107 | 0.3659 | 0.3301 | 0.1390 | 0.3454 | 0.9478 |
| 36 | 0.0063 | 0.0942 | 0.0902 | 0.0448 | 0.1386 | 0.4346 |
| 37 | 0.0714 | 0.3635 | 0.6713 | 0.1368 | 0.6630 | 0.5061 |
| 38 | 0.0322 | 0.2629 | 0.4181 | 0.0978 | 0.4049 | 0.6664 |
| 39 | 0.0001 | 0.0299 | 0.0102 | 0.0317 | 0.0745 | 0.3268 |
| 40 | 0.0143 | 0.1643 | 0.0933 | 0.1299 | 0.2784 | 0.3395 |
| 41 | 0.0023 | 0.0562 | 0.0371 | 0.0375 | 0.0977 | 0.5106 |

### 3.3. Comparison of Different Risk Stratification Methods

The performance of different risk stratification methods (Section 2.6) is assessed using the “ground-truth” risk data of 41 ATAA patients in Group 2. For Method 1∼4, FP of the 41 patients were computed using the patient-specific geometries and systolic blood pressures (see Table 3). Using the maximum diameter criterion (Method 1: *p* = 0.1677), no statistical significance was found between the high and low risk groups. Using the maximum FP evaluated at *P*_*sys*_, we found a significant difference between the high and low risk groups (Method 2: *p* = 0.0117). Difference between high and low risk groups is also significant (Method 3: *p* = 0.0070) when using representative hyperelastic properties and the maximum FP at 1.5*P*_*sys*_. Using patient-specific hyperelastic properties and the maximum FP at 1.5*P*_*sys*_, the lowest p-value is achieved (Method 4: *p* = 0.0001) to separate the high and low risk groups. The results are shown in Figure 7.

**Figure 7.**
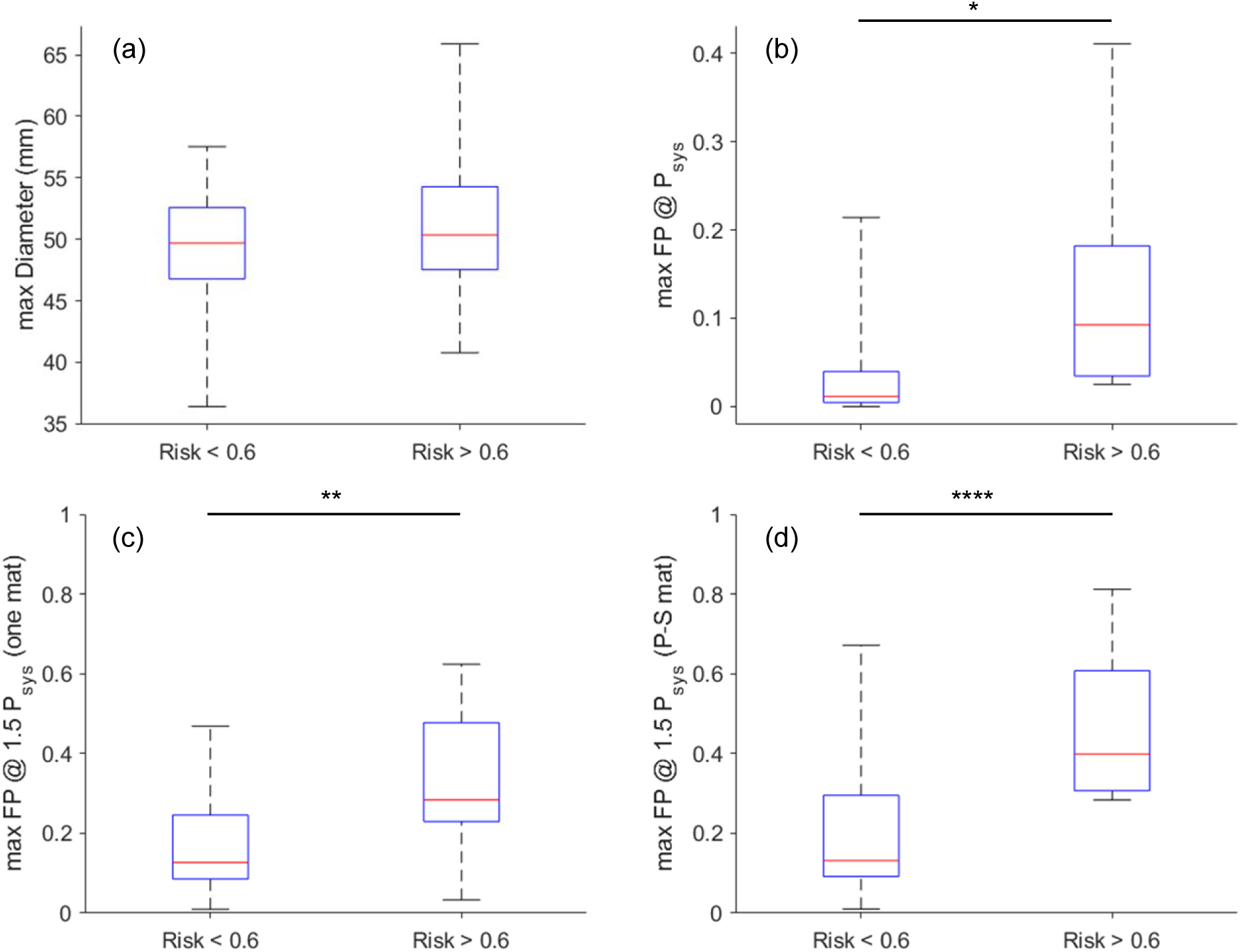
Distribution of failure metrics for high and low risk patients using different stratification methods (Section 2.6). The numerically-reconstructed PRR (Section 2.5) is used as the “ground-truth” risk displayed in the horizontal axis. (a) Method 1: maximum diameter; (b) Method 2: maximum FP evaluated at *P*_*sys*_; (c) Method 3: maximum FP evaluated at 1.5 *P*_*sys*_ using representative hyperelastic properties; and (d) Method 4: maximum FP evaluated at 1.5*P*_*sys*_ using patient-specific hyperelastic properties. “one mat” stands for representative hyperelastic properties; “P-S mat” stands for patient-specific hyperelastic properties. The red mark indicates the median, and the bottom and top edges of the box indicate the 25th and 75th percentiles, respectively. The whiskers extend to the maximum and minimum. *, **, **** indicates statistical significance levels of *p ≤* 0.05, *p ≤* 0.01, *p ≤* 0.0001, respectively.

ROC curves of the Method 1∼ Method 4 are shown in Figure 8. The areas under the curves (AUC), which reflect the discriminative powers of the failure metrics, are 0.5489, 0.8448, 0.7644 and 0.8621, respectively, for Method 1∼ Method 4. The diameter criterion has the lowest AUC, while the highest AUC is achieved by FP evaluated under elevated blood pressure using the patient-specific hyperelastic properties, which highlights the potential benefits of incorporating patient-specific hyperelastic properties [53-58] in the risk stratification. Comparing to FP at *P*_*sys*_ without patient-specific hyperelastic properties, the performance is not improved by evaluating FP under elevated blood pressure by using a representative set of hyperelastic parameters.

**Figure 8.**
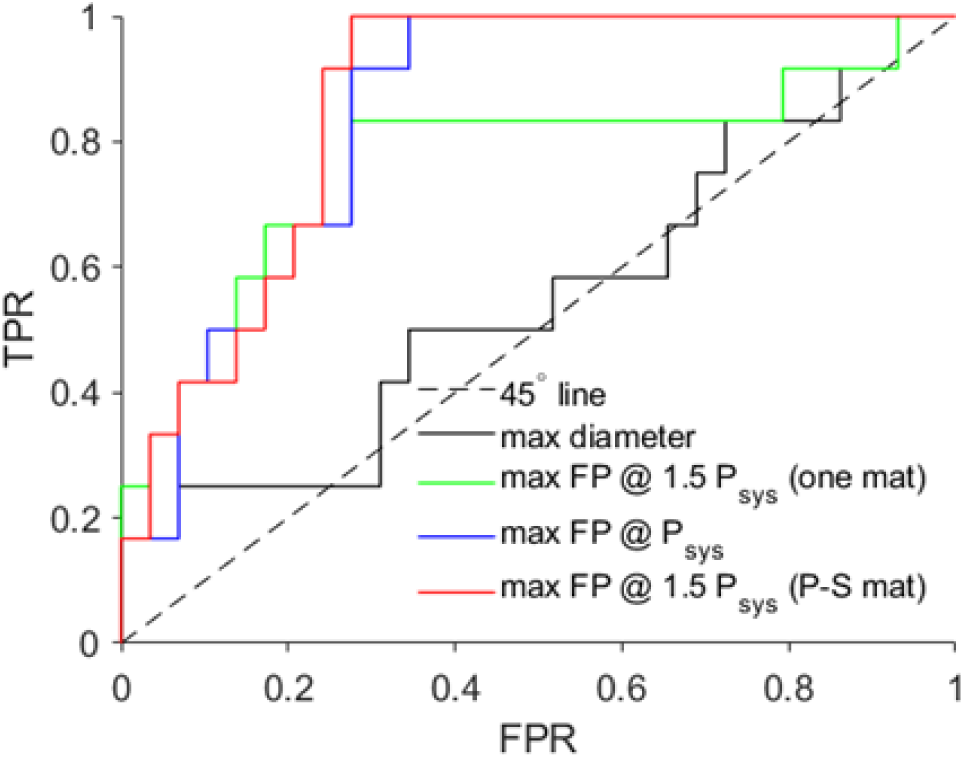
ROC curves of different risk stratification methods. The plots are generated using false positive rate (FPR) versus true positive rate (TPR). “one mat” stands for representative hyperelastic properties; “P-S mat” stands for patient-specific hyperelastic properties. AUC for the diameter (Method 1), FP at *P*_*sys*_ (Method 2), FP at 1.5*P*_*sys*_ using representative hyperelastic properties (Method 3), and FP at 1.5*P*_*sys*_ using patient-specific hyperelastic properties (Method 4) are 0.5489, 0.8448, 0.7644 and 0.8621, respectively.

In general, to evaluate a diagnostic method, an AUC of 0.5 suggests no discrimination, 0.7 to 0.8 is considered acceptable, and 0.8 to 0.9 is considered excellent [59].

### 3.4. Comparison between Probabilistic and Deterministic metrics

Method 2 and Method 4 (Section 2.6) can be equipped with the probabilistic metric (FP in Eqn. (11)) or the deterministic Tsai−Hill (TH) metric (*Λ* in Eqn. (2) with typical failure parameters *X* = 2.5*MPa* and *Y* = 1.2*MPa)*. FP and TH of the 41 ATAA patients (Group 2) at *P*_*sys*_ and 1.5*P*_*sys*_ were computed by using the patient-specific geometries and systolic blood pressures. The results are shown in Table 3. Difference between high and low risk groups is significant *(p* = 0.0099) using the maximum TH *at P*_*sys*_ (Method 2), while *p* = 0.0121 when the same method is used with FP. Using patient-specific hyperelastic properties and the maximum TH at 1.5*P*_*sys*_ (Method 4), the p-value is low *(p* = 0.0017). The most significant difference was found for FP using Method 4 (p = 0.0001). The results are summarized in Figure 9.

**Figure 9.**
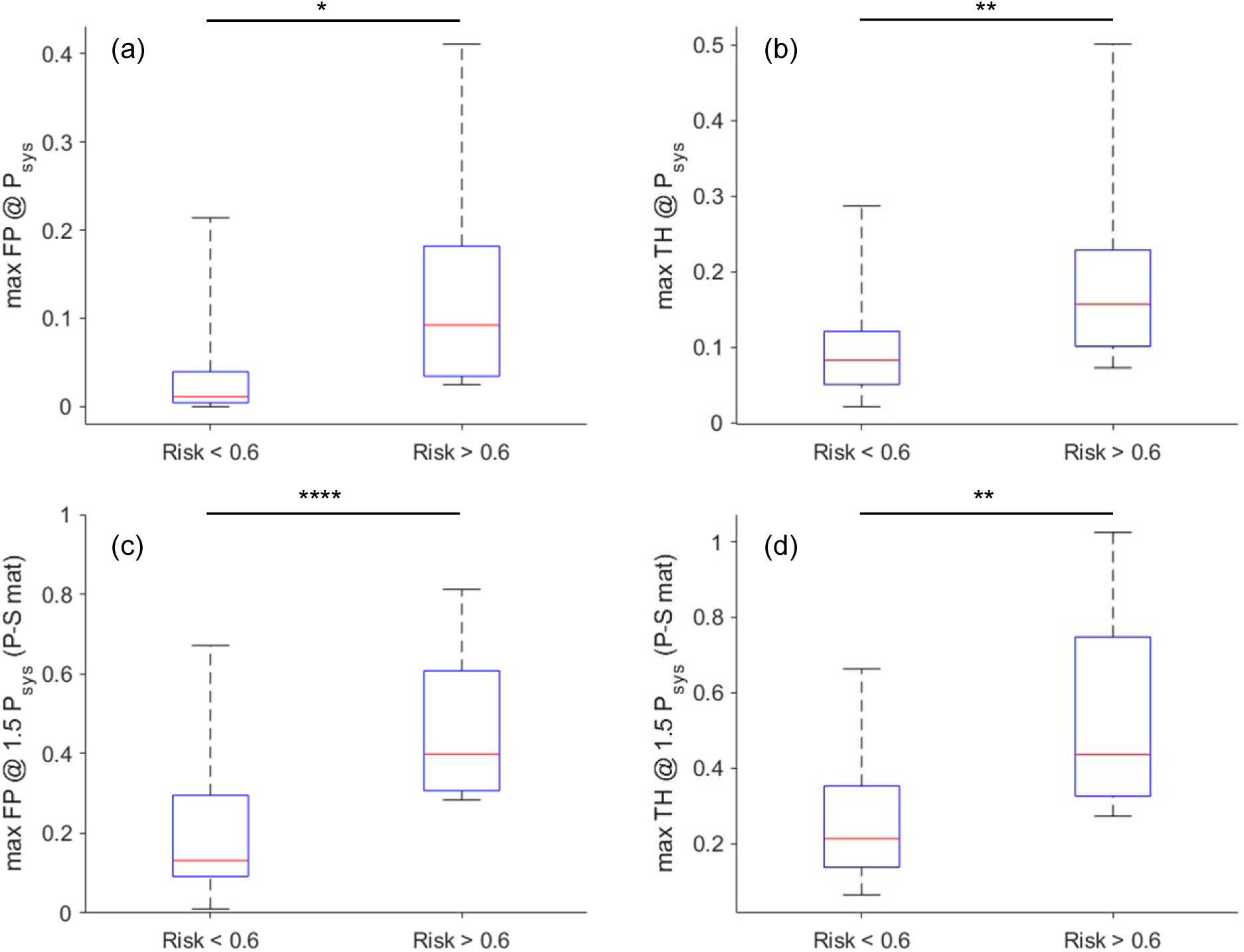
Distribution of probabilistic (FP) and deterministic (TH) failure metrics for high and low risk ATAA. The numerically-reconstructed PRR (Section 2.5) is used as the “ground-truth” risk displayed in the horizontal axis. (a) and (b) Method 2: FP and TH evaluated at *P*_*sys*_; (c) and (d) Method 4: FP and TH evaluated at 1.5*P*_*sys*_ using patient-specific hyperelastic properties. “P-S mat” stands for patient-specific hyperelastic properties. The red mark indicates the median, and the bottom and top edges of the box indicate the 25th and 75th percentiles, respectively. The whiskers extend to the maximum and minimum. *, **, **** indicates statistical significance levels of *p* ≤ 0.05, *p* ≤ 0.01, *p* ≤ 0.0001, respectively.

ROC curves of the maximum FP and the maximum TH metrics are demonstrated in

Figure 10 for failure metrics evaluated at *P*_*sys*_ (Figure 10(a)) and 1.5*P*_*sys*_ (Figure 10(b), using patient-specific hyperelastic properties). The AUC is 0.8448, 0.8017, 0.8621 and 0.8362, respectively, for FP at *P*_*sys*_, TH at *P*_*sys*_, FP at 1.5*P*_*sys*_ and TH at 1.5*P*_*sys*_. The results indicate that the probabilistic metric FP has a better discriminative power comparing to the deterministic metric TH.

**Figure 10.**
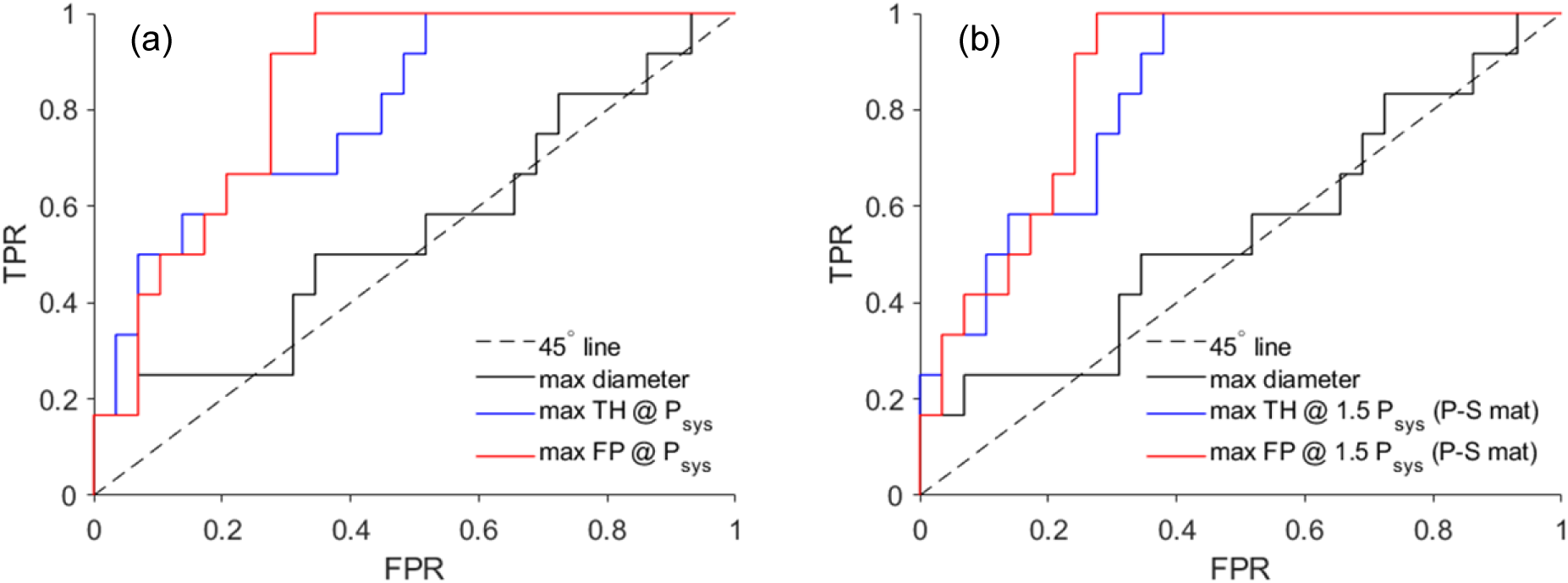
ROC curves of diameter criterion, probabilistic (FP) and deterministic (TH) failure metrics. The plots are generated using false positive rate (FPR) versus true positive rate (TPR). (a) Method 2: FP and TH evaluated at *P*_*sys*_; (b) Method 4: FP and TH evaluated at 1.5P_sys_ using patient-specific hyperelastic properties. “P-S mat” stands for patient-specific hyperelastic properties. AUC for the diameter, FP at *P*_*sys*_, TH at *P*_*sys*_, FP at 1.5*P*_*sys*_ and TH at 1.5*P*_*sys*_ are 0.5489, 0.8448, 0.8017, 0.8621 and 0.8362, respectively.

## 4. Discussion

In this study, a probabilistic anisotropic failure metric (FP) is proposed for ATAA risk stratification. Significant anisotropic failure properties of ATAA tissues have been shown in several studies [14, 15, 18-20], consistent with the testing results obtained in this study (see Figure 6). By using the TH criterion, anisotropic failure properties are embedded in FP. Comparing to the standard TH metric, FP incorporates uncertainties of wall strengths, i.e., the distribution of the failure parameters *X* and *Y* in the ATAA population. With more tissue failure testing data collected, FP can be updated and improved, e.g., a conditional FP can be built based on patient-specific information (age, gender, family history). Using reconstrued data, this study demonstrated that the performance of FP is superior to the standard TH metric with typical failure parameters. To our best knowledge, this is the first study that develops a probabilistic and anisotropic failure metric for quantifying failure risk of the aortic wall.

Using the reconstructed risk of 41 ATAA patients, we found that the biomechanical failure metrics have more discriminative power (AUC) than the maximum diameter criterion, which is consistent with recent studies [6, 7] investigating real ruptured and intact cases of abdominal aortic aneurysm (AAA). Using patient-specific hyperelastic properties and FP, the highest AUC is achieved (Figure 6), which highlights the potential benefit of identifying patientspecific hyperelastic properties [53-58] for risk stratifications. When patient-specific hyperelastic properties are not available, no additional benefit was found by assuming representative properties. In this case, failure metric can be evaluated on image-derived geometries using static determinacy [50-52] without the need of patient-specific hyperelastic properties. In this study, an ideal scenario was investigated, in which the wall thickness was assumed to be accurately measured from MRI [22-25] or high-resolution CT scans [26]. Hence, AUCs of the failure metrics could be close to the upper limits. Opposite to the promising results, a retrospective study [60] suggested no added value of biomechanical indices for AAA risk assessment. However, in their biomechanical model, the imaged-derived configuration was directly pressurized without proper consideration of zero-pressure configuration and pre-stress, which may result in inaccurate stress calculation.

Polzer and Gasser [26] proposed a probabilistic rupture risk index (PRRI) based on isotropic wall strength leveraging an uncertainty quantification (UQ) framework [61]. In their work [26], the wall thickness, which is the input to the FE simulations, was treated as a source of uncertainties. However, in the UQ framework, sampling-based approaches like Monte Carlo are typically needed to quantify the uncertainties of the FE output (i.e. peak wall stress) propagated from the uncertainties of the FE inputs (i.e., wall thickness and material parameters), which often results in high computational cost [62-64]. In contrast to PRRI [26] that incorporates uncertainties of both wall strength and wall thickness, the proposed probabilistic failure metric (FP) only considers uncertainties of anisotropic wall strength, and uncertainties from FE inputs are not involved. Therefore, given FE-computed stresses, FP can be derived from experimental uniaxial testing data via simple numerical integration. We have recently developed a deep learning (DL) model [61] as a fast and accurate surrogate of FE simulation, which can replicate the results of FE simulations instantaneously. To incorporate uncertainties originated from FE inputs such as wall thickness into FP, the DL model can be used to accelerate the UQ framework.

ATAA is a silent killer, the majority of patients remain asymptomatic until rupture or dissection occurs [1], which makes it difficult to obtain clinical CT images of ruptured ATAA. Sample size for ruptured AAA is also a challenge [65], for instance, a ruptured sample size of 7 was used by Polzer and Gasser [26]. A prospective study [6] collected 13 ruptured cases in about 2∼3 years of follow up period. In this study, ATAA risks (PRR) were retrospectively reconstructed by FE simulation using pre-operative images and tissue testing data of 41 patients who underwent elective surgery. These patient-specific images and tissue sample are often more accessible from clinics. We may be able to expand our validation data size once more CT images and tissue samples are collected.

In the current work, reconstruction of ATAA risk (PRR) was based on the following two assumptions: (1) Wall thickness and material properties were assumed to be homogenous across the ATAA geometry. It is known that the hyperelastic and failure properties of aortic wall are regional dependent [18, 66]. The wall thickness may also have spatial variations, and heterogeneity of wall thickness and material heterogeneity could be correlated [67]. In the FE simulations, a simplified case was considered, where the wall thickness was calculated from an averaged experimentally measured value, and the experimentally-derived hyperelastic behavior was used for the entire ATAA geometry. (2) Residual stresses were not incorporated in the FE simulations. In future studies, residual deformation could be taken into considerations by means of the GPA algorithm [65] or volumetric growth approach [68]. However, it is worth noting that the mean wall stress computed by Method 2 (Section 2.6) is independent of residual stresses [50].

## 5. Conclusions

In this work, we develop a novel probabilistic and anisotropic failure metric using failure property data from 84 ATAA patients. The novel failure probability (FP) metric, which was derived based on the Tsai−Hill (TH) failure criterion, incorporates uncertainties of the anisotropic failure properties. Numerically-reconstructed risks of additional 41 ATAA patients were used for validation. Performance of different risk stratification methods (diameter, with or without patient-specific hyperelastic properties) were compared. The probabilistic FP metric outperforms the deterministic TH metric using the reconstructed data. The results also revealed potential benefit of identifying patient-specific hyperelastic properties in ATAA risk stratification.

## 6. Acknowledgment

This study is supported by American Heart Association (AHA) 18TPA34230083. We thank previous and current lab members, Thuy Pham, Caitlin Martin, Fatiesa Sulejmani, M. Subhi Al Jabi, Parikirt Oggu, Ahdil Gill, and Clarisse Gieowarsingh for assistance in collecting the tissue mechanical testing data, Vishal Shah for assistance in data analysis, and Daksha Jadhav for assistance in image segmentation.

